# Multi-metrics assessment of dynamic cerebral autoregulation in middle and posterior cerebral arteries in young fit women

**DOI:** 10.1101/2020.05.25.114447

**Authors:** Lawrence Labrecque, Audrey Drapeau, Kevan Rahimaly, Sarah Imhoff, François Billaut, Patrice Brassard

## Abstract

Individuals with low orthostatic tolerance show greater decrease in posterior cerebral artery mean blood velocity (PCAv_mean_). Since young fit women often experience presyncopal symptoms, their posterior cerebral circulation may be prone to greater decreases in PCAv_mean_, probably explained by an attenuated dynamic cerebral autoregulation (dCA). Regional differences in dCA have never been evaluated in young fit women. We compared dCA in the middle cerebral artery (MCA) and posterior cerebral artery (PCA) in 11 young fit women (25 ± 4y; 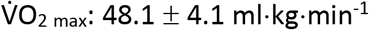) in response to a sit-to-stand (5 min sitting followed by 5 min standing) and repeated squat-stand maneuvers performed at 0.05 Hz and 0.10 Hz. The cerebral pressure-flow relationship was characterized using four metrics: 1) percent reduction in blood velocity (BV) per percent reduction in MAP (% BV/% MAP) during initial orthostatic stress (0-15 s after sit-to-stand); 2) onset of the regulatory response (i.e. time delay before an increase in conductance (BV/MAP); 3) rate of regulation (RoR), following sit-to-stand and; 4) transfer function analysis (TFA) of forced MAP oscillations induced by repeated squat-stands. Upon standing, the relative decline in MCAv_mean_ and PCAv_mean_ was similar (−25 ± 9 vs. −30 ± 13%; p=0.29). The onset of the regulatory response (p=0.665), %ΔBV/%ΔMAP (p=0.129) and RoR (p=0.067) were not different between MCA and PCA. In regard to TFA, there was an ANOVA artery effect for gain (p<0.001) and a frequency effect for phase (p<0.001). These findings indicate the absence of regional differences in dCA in young fit women.

**New findings:** *What is the central question of this study?:* Are there regional differences in the dynamic cerebral autoregulation in young fit women?

*What is the main finding and its importance?:* The key finding of this study is that there are no differences in dynamic cerebral autoregulation between both arteries. These results indicate that dynamic cerebral autoregulation does not seem to be responsible for making the posterior cerebral circulation more vulnerable to transient reduction in blood pressure in young fit women.

## Introduction

Syncope is associated with a decrease in cerebral blood velocity (CBV), often monitored in middle cerebral arteries (MCA) (Kay, Sprick, & Rickards, 2017; Schondorf, Benoit, & Stein, 2001; van Lieshout, Wieling, Karemaker, & Secher, 2003). However, both MCAs supply the anterior part of the brain and questions arise as to whether the posterior irrigation could rather be linked to the appearance of pre-syncopal symptoms. Vertebral arteries and their branches supply blood to cardiac, vasomotor and respiratory centers in the medulla oblongata (Tatu, Moulin, Vuillier, & Bogousslavsky, 2012). Emergence of orthostatic symptoms (such as dizziness, nausea and blurred vision) could therefore refer to posterior cerebral hypoperfusion, notably in posterior cerebral arteries (PCA). Kay & Rickards (2016) demonstrated that individuals with low tolerance to orthostatic stimulation show greater decrease in PCA mean blood velocity (PCAv_mean_) during a progressive lower body negative pressure protocol where tolerance was evaluated as the onset of presyncope/appearance of symptoms (Kay et al., 2017). Conversely, these investigators did not report any differences in MCA mean blood velocity (MCAv_mean_) changes during the protocol compared to baseline. Of note, young women display greater prevalence of orthostatic intolerance (Fu et al., 2004) and symptoms of cerebral hypoperfusion than young men (Ali et al., 2000). Therefore, their posterior cerebral circulation could be prone to greater decreases in blood flow when blood pressure lowers (Kay et al., 2017).

An attenuated dynamic cerebral autoregulation (dCA), which describes the ability of the cerebrovasculature to respond to rapid changes in mean arterial pressure (MAP), in the posterior cerebral circulation could accentuate the reduction in blood supply to posterior regions of the brain following acute MAP reduction, which in turn could explain the appearance of symptoms in women during orthostatic stress. However, although we previously reported that healthy young fit women display diminished dCA in the MCA compared to men (Labrecque et al., 2019a), whether a differential dCA response between MCA and PCA in these young fit women remains to be clearly determined.

Few studies have examined dCA in anterior compared to posterior cerebral regions. Some findings, but not all, suggest regional differences in dCA. Sorond and coworkers did not report any differences between MCAv_mean_ and PCAv_mean_ responses following a sit-to-stand maneuver in young individuals (Sorond, Khavari, Serrador, & Lipsitz, 2005). Transfer function analysis (TFA) of spontaneous or forced oscillations in MAP and CBV can also be used to characterize dCA using diverse metrics (coherence: fraction of the MAP linearly related to CBV; gain: amplitude of CBV change for a given MAP change; and phase: difference in the timing of the MAP and CBV oscillations). Using repeated squat-stand maneuvers to force MAP oscillations of large amplitude, no regional differences in dCA metrics were reported in young healthy participants (Burma et al., 2020; Smirl, Hoffman, Tzeng, Hansen, & Ainslie, 2015). Conversely, Haubrich et al. described higher PCA TFA gain of spontaneous oscillations in both supine position and following head-up tilt (HUT), which has been interpreted by these authors as a lowered dampening capacity of the PCA (Haubrich, Wendt, Diehl, & Klotzsch, 2004). Interestingly, Wang et al. reported a greater relative decline in PCAv_mean_ compared to MCAv_mean_ following HUT, but only in women (Yuh-Jen Wang, 2010). This latter observation highlights the importance of studying dCA in anterior and posterior cerebral circulation specifically in women.

The equivocal results from these studies examining dCA in both MCA and PCA could be explained, at least in part, by the various methodologies used to assess dCA. Analysis of spontaneous oscillations is controversial since it is not associated with high coherence (Smirl et al., 2015). In fact, low coherence may suggest low signal-to-noise ratio and it cannot be assumed that MAP and CBV signals are linearly correlated. Also, TFA of spontaneous oscillations has poor reproducibility. Even though the utilization of repeated squats-stands to force blood pressure oscillations is considered as the gold standard for linear TFA metric interpretation (Smirl et al., 2015), this maneuver may solicit the cardiorespiratory system to a significant degree for some individuals and may not represent a MAP challenge associated with activities from daily living such as a sit-to-stand maneuver.

With the understanding that each dCA assessment technique, including MAP stimuli of various natures and amplitudes, most likely reflects somewhat different components of the cerebral-pressure flow relationship (Labrecque et al., 2019a; 2017; Tzeng et al., 2012), the aim of this study was to compare dCA in the MCA and PCA in young healthy fit women using a multi-metrics approach, including 1) a sit-to-stand maneuver and 2) TFA analysis of forced oscillations in MAP and CBV induced by repeated squats-stands. We hypothesized PCA would show a diminished buffering capability of MAP changes.

## Methods

### Ethical Approval

All participants provided written informed consent prior to participating in the investigation, and the study was approved by the *Comité d’éthique de la recherche de l’Institut universitaire de cardiologie et de pneumologie de Québec – Université Laval* (CER: 21180) according to the principles established in the Declaration of Helsinki (except for registration in a database).

### Participants

Eleven moderately-trained women were recruited for this study (age: 25 ± 4 yrs, height: 1.64 ± 0.07 m, body mass: 61.0 ± 5.7 kg, body mass index: 23 ± 2 kg/m^2^, maximal oxygen uptake 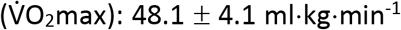, training volume: 465 ± 159 min/week). All the participants trained in a variety of endurance-based sports including cycling (n=1), triathlon (n=4), mountain biking (n= 1), running (n= 4) and cross-country skiing (n=1). All participants were healthy and free from any medical conditions, demonstrated a normal 12-lead electrocardiogram (ECG), and were not taking medications. Women were either taking oral contraceptive continuously since > 1 year (n=2), wearing an intrauterine device (n=2) or were tested during menses or the early follicular phase (day 1 to 10) of their menstrual cycle (n=7).

### Experimental protocol

This study was part of a larger study examining the influence of an elevated cardiorespiratory fitness on dCA in young healthy women and the cerebrovascular response to an acute bout of high-intensity exercise (Labrecque et al., 2019a; 2020). However, the current question was determined a priori and was prospectively studied as a separate question. Anthropometrics measurements, 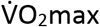 and cerebrovascular haemodynamics of the MCA have previously been published (Labrecque et al., 2019a; 2020). Participants visited the laboratory twice to perform: 1) an incremental cycling protocol for 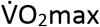 determination, and 2) anthropometrics, systemic and cerebrovascular resting measurements and dCA characterization. Participants were asked to avoid exercise training for a minimum of 12 h, as well as alcohol, drugs and caffeine consumption for 24 h before each visit. All visits and evaluations were performed in the same order for all participants and there was at least 48 h between each visit. During all the cerebrovascular experiments, participants were asked to keep their eyes opened and we ensured there was minimal visual stimuli.

### Measurements

#### Systemic haemodynamics

Heart rate (HR) was measured using a 5-lead ECG. Beat-to-beat BP was measured by the volume-clamp method using a finger cuff (Nexfin, Edwards Lifesciences, Irvine, CA, USA). The cuff was placed on the right middle finger and referenced to the level of the heart using a height correct unit for BP correction. MAP was obtained by integration of the pressure curve divided by the duration of the cardiac cycle. This method has been shown to be a reliable index of the dynamic changes in beat-to-beat BP which correlate well with the intra-arterial BP recordings and can be used to describe the dynamic relationship between BP and cerebral blood velocity (Omboni et al., 1993; Sammons et al., 2007).

#### Middle and posterior cerebral artery blood velocities

MCAv_mean_ and PCAv_mean_ were monitored with a 2-MHz pulsed transcranial Doppler ultrasound (Doppler Box; Compumedics DWL USA, Inc. San Juan Capistrano, CA). Identification and location of the left MCA and right PCA were determined using standardized procedures (Willie et al., 2011). Probes were attached to a headset and secured with a custom-made headband and adhesive conductive ultrasonic gel (Tensive, Parker Laboratory, Fairfield, NY, USA) to ensure a stable position and angle of the probe throughout testing.

#### End-tidal partial pressure of carbon dioxide

End-tidal partial pressure of carbon dioxide (P_ET_CO_2_) was continuously measured during all the evaluations through a breath-by-breath gas analyzer (Breezesuite, MedGraphics Corp., MN, USA; Ultima™ CardiO2^®^ Gas Exchange Analysis System, MGC Diagnostics^®^, MN, USA) calibrated to known gas concentrations following manufacturer instructions before each evaluation.

#### Data acquisition

For each assessment, signals (except for P_ET_CO_2_) were analog-to-digital converted at 1kHz via an analog-to-digital converter (Powerlab 16/30 ML880; ADInstruments, Colorado Springs, CO, USA) and stored for subsequent analysis using commercially available software (LabChart version 7.1; ADInstruments). P_ET_CO_2_ was time-aligned with the other signals.

### Visit 1

#### Maximal oxygen consumption 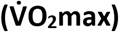

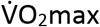 was determined during a progressive ramp exercise protocol performed on an electromagnetically braked upright cycle ergometer (Corival, Lode, the Netherlands) to characterize cardiorespiratory fitness of the participants as previously described (Drapeau et al., 2019; Labrecque et al., 2017; 2020).

### Visit 2

#### Anthropometric measurements and resting haemodynamics

Height and body mass were measured in each participant. Resting haemodynamic measurements included MAP (volume-clamp method using a finger cuff), heart rate (HR; ECG), MCAv_mean_ and PCAv_mean_, which were continuously monitored on a beat-by-beat basis during 5 min of seated rest. Cerebrovascular conductance index (CVCi; MCAv_mean_ or PCAv_mean_/MAP) and its reciprocal, resistance (CVRi; MAP/MCAv_mean_ or PCAv_mean_) were then calculated. P_ET_CO_2_ (gaz analyzer) was continuously monitored on a breath-by-breath basis. The average values of the last minute of recording represented the baseline.

### Assessment of dynamic cerebral autoregulation

#### Sit-to-stand

Following 5 min of seated rest, participants quickly (<2 s) stood up and remained in a standing position for 5 min without moving or contracting leg muscles. HR, MAP, MCAv_mean_, PCAv_mean_ and P_ET_CO_2_ were continuously monitored during the sit-to-stand procedure. The sit-to-stand had to cause a decrease in MAP > 10 mmHg to be considered valid and to be included in the analysis (Subudhi et al., 2015). Additionally, we evaluated the prevalence of initial orthostatic hypotension (IOH), defined as a decrease in systolic BP ≥ 40 mmHg and/or a decrease in diastolic BP ≥ 20 mmHg during the first 15 s of standing (Finucane et al., 2019; Freeman et al., 2011).

#### Forced blood pressure oscillations using repeated squat-stand maneuvers

Following a minimum of 10 min in a standing quiet rest to ensure stabilization of baseline haemodynamics following the sit-to-stand maneuver, repeated squat-stands were performed as previously described (Drapeau et al., 2019; Labrecque et al., 2019a). Briefly, participants were asked to alternate standing and squatting positions during 5 min at frequencies of 0.05 Hz (10-s squat, 10-s standing) and 0.10 Hz (5-s squat, 5-s standing). The sequence of the repeated squat-stands was randomized between participants and each frequency was separated by 5 min of standing recovery. During these maneuvers, participants were instructed to maintain normal breathing and to avoid Valsalva. The linear aspect of the dynamic MAP-MCAv/PCAv relationship was characterized via TFA (see the “Data analysis and statistical approach” section). MAP, HR, MCAv, PCAv and P_ET_CO_2_ were continuously monitored during this evaluation. An averaged P_ET_CO_2_ of the first and last five breaths of each maneuver (0.05 and 0.10 Hz) was calculated.

### dCA calculations

#### Acute cerebrovascular responses to acute reduction in blood pressure induced by a sit-to-stand

The following metrics were used to characterize the cerebral pressure-flow relationship to acute reduction in blood pressure induced by the sit-to-stand maneuver: 1) the reduction in MAP, MCAv_mean_ and PCAv_mean_ to their respective nadir (absolute: ΔMCAv_mean_, ΔPCAv_mean_, ΔMAP; and relative to baseline: ΔMCAv_mean_ (%), ΔPCAv_mean_ (%), ΔMAP (%)); 2) the percent reduction in blood velocity (BV) per percent reduction in MAP (%ΔBV/%ΔMAP); 3) the time delay before the onset of the regulatory response; 4) the rate of decline in MCAv_mean_ and PCAv_mean_ and; 5) the rate of regulation (RoR).

1. The reduction in MAP and MCAv_mean_ or PCAv_mean_ is the difference between baseline MAP, MCAv_mean_ or PCAv_mean_ (averaged over the last 15 s of seated rest before standing) and minimum MAP, MCAv_mean_ or PCAv_mean_ recorded after the sit-to-stand.
2. %ΔBV/%ΔMAP upon standing was calculated for both MCA and PCA as follows: [((baseline BV – minimum BV)/baseline BV)/((baseline MAP – minimum MAP)/baseline MAP)].
3. The time delay before the onset of the regulatory response is the time lapse between the beginning of the sit-to-stand and the elevation in CVCi (18). The onset of the regulatory response becomes visible when CVCi begins to continuously increase (without any subsequent transient reduction) during acute reduction in blood pressure. This metric was assessed by two different observers (LL and PB).
4. The rate of decline in MCAv_mean_ and PCAv_mean_ upon standing was calculated as follows: [(%MCAv_mean_ (or %PCAv_mean_) at nadir - %MCAv_mean_ (or %PCAv_mean_) before decline)/(Δt)].
5. The physiological response to an acute reduction in blood pressure can be divided into two phases (Ogoh, Brothers, Eubank, & Raven, 2008); Phase I is the time point after the sit-to-stand where MCAv_mean_ or PCAv_mean_ changes are independent of any arterial baroreflex correction (1 to 7 s after sit-to-stand) (Deegan, Sorond, Lipsitz, ÓLaighin, & Serrador, 2009; Sorond, Serrador, Jones, Shaffer, & Lipsitz, 2009; van Beek, Claassen, Rikkert, & Jansen, 2008). Phase II is the time point starting at the onset of arterial baroreflex and continuing for 4 s (Ogoh et al., 2008). During Phase I, the rate of change in CVCi is directly related to dCA, without arterial baroreflex regulation (Aaslid, Lindegaard, Sorteberg, & Nornes, 1989). RoR was calculated during Phase I using the following equation:

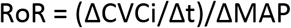 Where ΔCVCi/Δt is the linear regression slope between CVCi and time (t) during Phase I (a 2.5-s interval (Δt) after individually determined onset of the regulatory change following sit-to-stand was used for the analysis of RoR), and ΔMAP is calculated by subtracting baseline MAP from averaged MAP during Phase I (Aaslid et al., 1989; Ogoh et al., 2008).

#### Assessment of the dynamic relationship between MAP and MCAv or PCAv

Data from forced blood pressure oscillations were analyzed using commercially available software Ensemble (Version 1.0.0.14, Elucimed, Wellington, New Zealand) in accordance with the recommendations of the Cerebral Autoregulation Research Network (CARNet) (Claassen, Meel-van den Abeelen, Simpson, Panerai, international Cerebral Autoregulation Research Network (CARNet), 2016). Beat-to-beat MAP and MCAv or PCAv signals were spline interpolated and re-sampled at 4 Hz for spectral analysis and TFA based on the Welch algorithm. Each 5-min recording was first subdivided into 5 successive windows that overlapped by 50%. Data within each window were linearly detrended and passed through a Hanning window prior to discrete Fourier transform analysis. For TFA, the cross-spectrum between MAP and MCAv or PCAv was determined and divided by the MAP auto-spectrum to derive the transfer function coherence, absolute gain (cm/s/mmHg), normalized gain (nGain; %/mmHg) and phase (radians). TFA coherence, gain and phase of forced MAP oscillations were sampled at the point estimate of the driven frequency (0.05 and 0.10 Hz). These point estimates were selected as they are in the very low (0.02-0.07 Hz) and low (0.07-0.20 Hz) frequency ranges where dCA is thought to be most operant (Smirl et al., 2015). Only the TFA phase and gain values where coherence exceeded 0.50 were included in analysis to ensure the measures were robust for subsequent analysis. Phase wrap-around was not present at any of the point-estimate values for repeated squat-stands.

#### Statistical analysis

The normal distribution of data was confirmed using Shapiro-Wilk normality tests. Baseline differences and sit-to-stand related metrics between MCAv_mean_ and PCAv_mean_ were analyzed with paired t-tests. Wilcoxon tests were used if data were not normally distributed. Two-way ANOVAs (factors: artery and frequency) were performed for each TFA metrics. Relationships between variables were determined using Pearson product-moment or Spearman’s Rho correlations. Statistical significance was established *a priori* at p <0.05 for all two-tailed tests. Data are expressed as mean ± standard deviation.

## Results

Two participants were excluded from the sit-to-stand analysis since MAP did not decrease >10mmHg. Two participants were excluded from all TFA analysis of forced MAP oscillations since they did not complete the evaluation and we were unable to monitor PCAv for one more participant. The final sample size was n=9 for the sit-to-stand maneuver, n=9 for TFA of forced MCAv oscillations and n=8 for TFA of forced PCAv oscillations. Baseline systemic and cerebrovascular haemodynamics for this group of young fit women are presented in Table 1. CBV and CVCi were greater in the MCA compared to PCA (both p <0.0001).

**Table 1.** Resting seated values

| **Resting values** |  |
| --- | --- |
| n | 11 |
| Heart rate (bpm) | 73 ± 11 |
| Mean arterial pressure (mmHg) | 96 ± 10 |
| Middle cerebral artery mean blood velocity (cm·s<sup>-1</sup>) | 75 ± 7 |
| Posterior cerebral artery mean blood velocity (cm·s<sup>-1</sup>) | 43 ± 7 |
| MCA cerebrovascular conductance index (cm·s<sup>-1</sup>·mmHg<sup>-1</sup>) | 0.78 ± 0.10 |
| PCA cerebrovascular conductance index (cm·s<sup>-1</sup>·mmHg<sup>-1</sup>) | 0.45 ± 0.09 |
| End-tidal partial pressure of carbon dioxide (mmHg) | 37 ± 2 |

Data are presented as mean ± SD.

### Sit-to-stand

Upon standing, MAP decreased by 24 ± 4 mmHg (26 ± 3%) in 8 ± 3 s. In the 9 women who presented a valid MAP decrease (> 10 mmHg), four reached the criteria for IOH. The absolute decrease in blood velocity was greater in MCA vs. PCA (−19.5 ± 7.8 vs. −12.5 ± 5.7 cm/s; p=0.002) whereas the relative decline was similar (−25 ± 9 vs. −30 ± 13%; p=0.29). The rate of decline in MCAv_mean_ and PCAv_mean_ (−5.21 ± 1.77 vs. −3.56 ± 3.29 %/s; p=0.21) and the time to reach their respective nadir (8 ± 3 s for both MCAv_mean_ and PCAv_mean_; p=0.75) were comparable between arteries (Figure 1). The onset of the regulatory response (p=0.665), %ΔBV/%ΔMAP (p=0.129) and RoR (p=0.067) were not different between MCA and PCA (Figure 2). No correlations were found between reductions in MCAv_mean_ or PCAv_mean_ and sit-to-stand metrics. For the MCA, there was a positive correlation between %ΔBV/%ΔMAP and TFA gain (r=0.81; p=0.022) and nGain (r=0.85; p=0.011) during 0.05 Hz squat-stands. There were no other correlations between sit-to-stand and TFA metrics.

**Figure 1.**
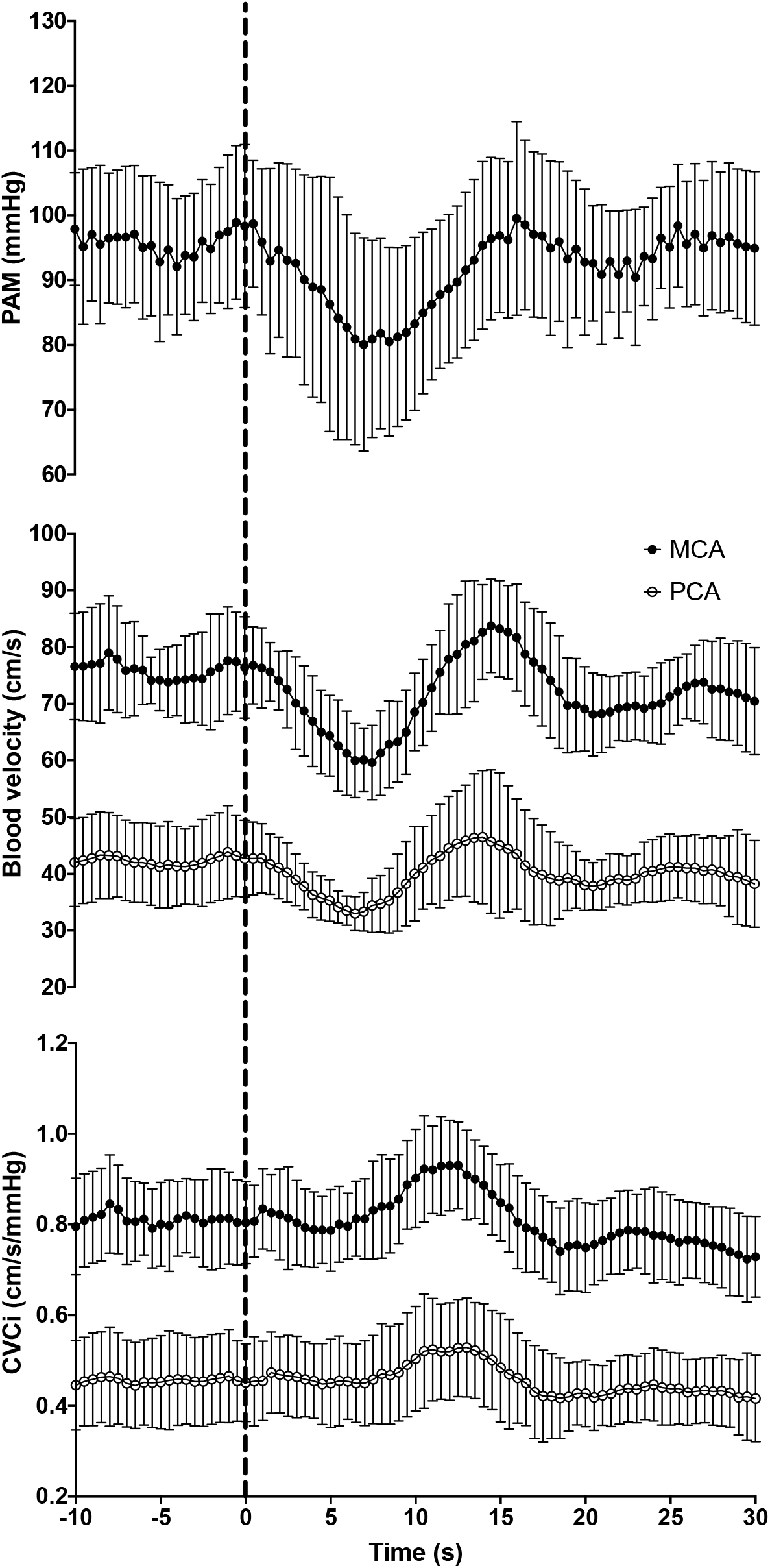
Mean arterial pressure (MAP), middle (MCA) and posterior cerebral arteries (PCA) mean blood velocity and cerebrovascular conductance indexes (CVCi) during sit-to-stand. Time 0 (dashed line) indicates the transition from the sitting to standing.

**Figure 2.**
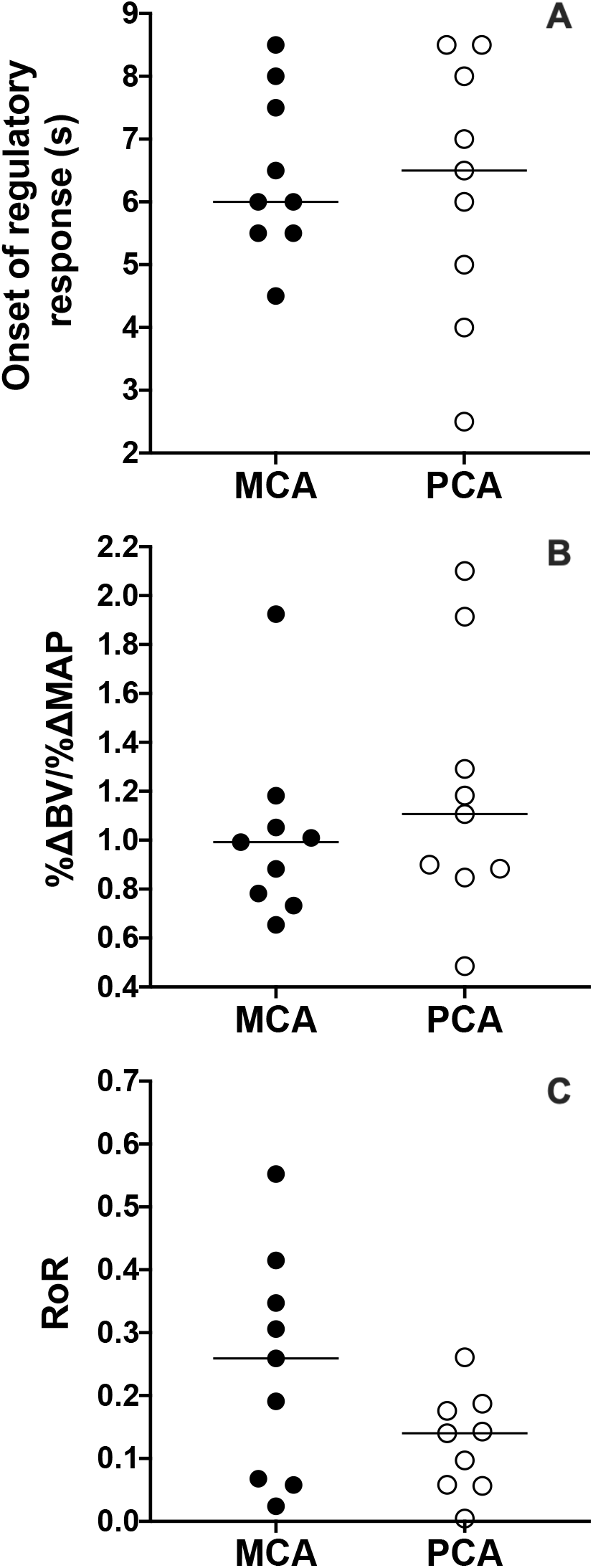
Cerebrovascular responses following sit-to-stand. Onset of regulatory response (A), percent reduction in blood velocity (BV) per percent reduction in MAP (%ΔBV/%ΔMAP; B) and the rate of regulatory response (C). Black circles indicate values for middle cerebral artery (MCA) and white circles indicates values for posterior cerebral artery (PCA).

### TFA of forced oscillations in MAP and MCAv/PCAv

MAP power spectrum densities during 0.05 Hz and 0.10 Hz repeated squat-stands were 2491 ± 2539 and 8517 ± 3424 mmHg^2^, respectively. CBV power spectrum density was greater in the MCA during 0.10 Hz (6504 ± 4161 vs. 3159 ± 2818 cm/s^2^; p=0.0027) but not 0.05 Hz (2180 ± 4321 vs. 987 ± 1832 cm/s^2^; p=0.1924) repeated squat-stands. There was an ANOVA artery effect for gain (p<0.001) and a frequency effect for phase (p < 0.001; Figure 3). Coherence and nGain were not different between arteries (Figure 3). The averaged P_ET_CO_2_ during each 5-min repeated squat-stands were not different between frequencies (0.05 Hz: 36.1 ± 3.3 vs. 0.10 Hz: 37.1 ± 2.8 mmHg; p= 0.08). Changes in P_ET_CO_2_ from the beginning to the end of repeated squat-stands executed at each frequency were also similar (0.05 Hz: +1.6 ± 1.2 vs. 0.10 Hz: + 1.4 ± 2.3 mmHg; p=0.50).

**Figure 3.**
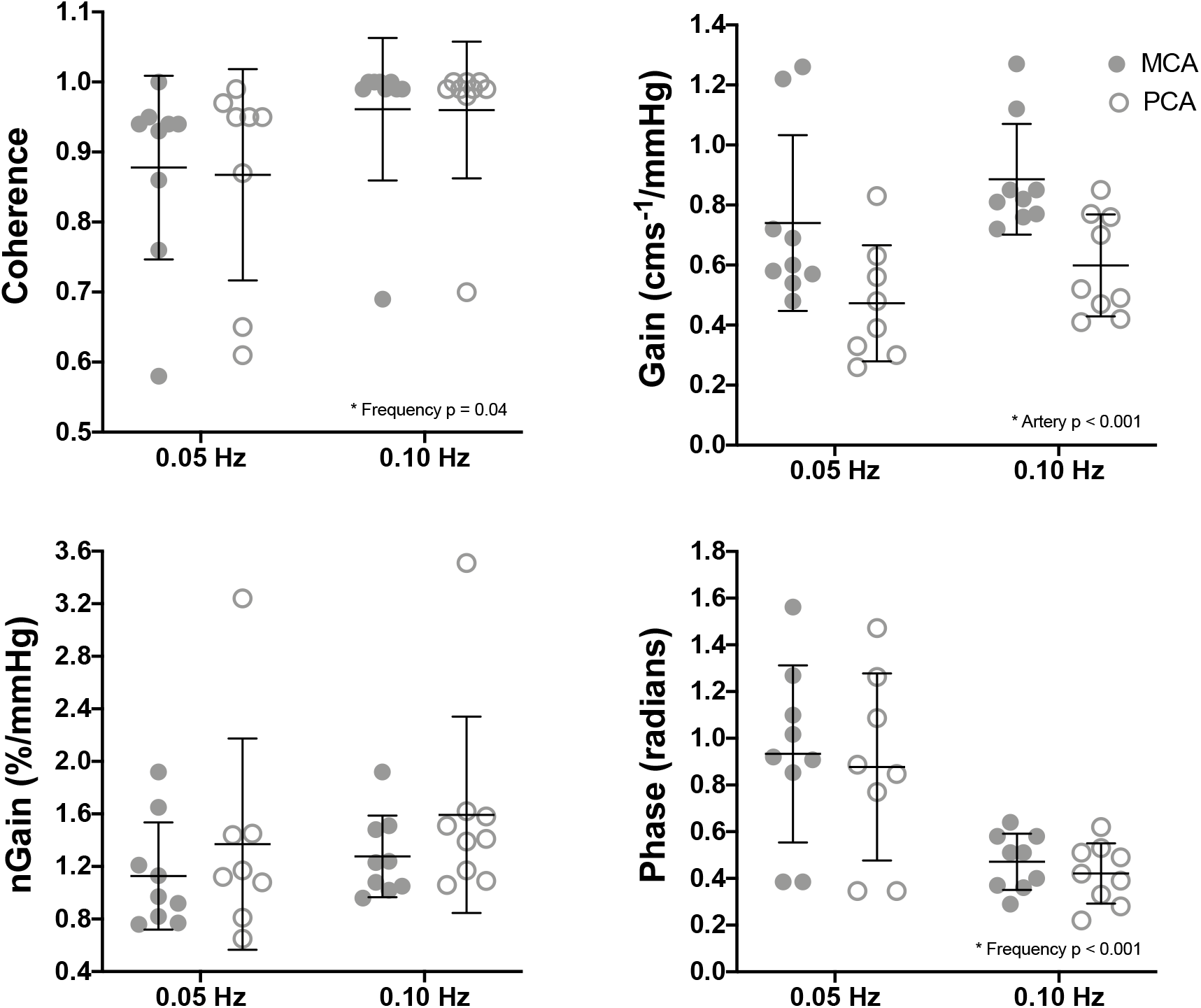
Transfer function analysis of forced oscillations in mean arterial pression and middle (MCA) and posterior cerebral artery (PCA) blood velocities. Group average coherence, gain, normalized gain (nGain) and phase for 0.05 Hz and 0.10 Hz squat-stands. Data are mean ± SD.

## Discussion

To the best of the authors’ knowledge, this is the first study to compare dCA in the MCA and PCA in young fit women using a multi-metrics approach. The key finding of this study is that there are no differences in dCA between both arteries. These results indicate that dCA might not be responsible for making the posterior cerebral circulation more vulnerable to transient reduction in blood pressure in young fit women.

### Resting cerebral haemodynamics

In healthy individuals, MCAv_mean_ is higher than PCAv_mean_ at rest (Burma et al., 2020; Haubrich et al., 2004; Sorond et al., 2005), which is concordant with our findings. The main reason for this difference resides in the MCAs being supplied by internal carotid arteries, which provide 75-80% of total cerebral blood flow, whereas PCAs are supplied by vertebral arteries, representing only 25-30% of the blood input to the brain (Sato et al., 2012).

### Acute reduction in blood pressure induced by a sit-to-stand

To the best of our knowledge, only one other study examined both MCA and PCA blood velocity responses during a sit-to-stand protocol, where investigators compared young and old participants (Sorond et al., 2005). No regional differences were identified in the young group. The authors observed comparable relative reductions in MAP, MCAv_mean_ and PCAv_mean_, as reported in the current study. Although not different between vessels, their RoR values were slightly higher than the RoR values from our participants for both vessels. It is important to note, however, that Sorond et al. (2005) used the method described by Aaslid (1989), based on the baroreflex onset with the time window from 0 to 2.5 s following thigh cuff release (Aaslid et al., 1989). We rather used the onset of the regulatory response (i.e. the moment when CBV starts to recover) as the beginning of our time interval for analysis (Labrecque et al., 2017; 2019a; Lind-Holst et al., 2011). The latest approach is more factual since in the first 2.5 s after standing, neither MAP or CBV have completed their reduction (Labrecque et al., 2019a). Interestingly, in older participants, Sorond et al. (2005) observed a greater PCAv_mean_ decline and a smaller reduction in cerebrovascular resistances, suggesting a vulnerability of posterior cerebral regions to hypoperfusion during acute reduction in blood pressure with advancing age (Sorond et al., 2005). Importantly, the inclusion of both men and women in the group composed of young participants could have influenced their results. Indeed, the literature remains equivocal in regards to sex differences in dCA (Favre & Serrador, 2019; Labrecque et al., 2019a) including a potential confounding effect of cardiorespiratory fitness (Labrecque, Smirl, & Brassard, 2019b). Additional studies using a transition to the standing position in order to reduce MAP have been published in children and adolescents. Following a supine-to-stand maneuver in adolescents, Vavilala & colleagues reported higher autoregulation index (ARI) in the MCA in boys and contrastingly, higher ARI in the basilar artery in girls (Vavilala et al., 2005). Several methodological issues make the comparison with our current findings difficult, including the fact that participants in the study by Vavilala et al. (2005) were children, that cerebral blood flow was assessed in the basilar artery, and the lack of details in regard to potential influence of visual stimuli.

In response to HUT, Wang et al. (2010) reported a greater decline in PCAv_mean_ compared to MCAv_mean_ in women only (Wang, Chao, Chung, Huang, & Hu, 2010). That observation could be the consequence of a diminished CA in the posterior circulation in women. However, the contrasting results between the study by Wang et al. and ours may be explained by a different initial body position before the transition to the standing position. There are in fact evidence that dCA is improved in the supine position when compared to seated or standing positions (Favre, Lim, Falvo, & Serrador, 2020). Therefore, the transition from supine to standing might cause greater cardiovascular and cerebrovascular challenges than the transition from sitting to standing. Interestingly, extracranial blood flow presents regional differences in response to HUT. Sato et al. reported that VA blood flow is maintained compared to ICA blood flow (Sato et al., 2012). However, when a thigh-cuff test is superimposed to HUT, changes in VA blood flow are greater and indices of dCA attenuated compared to the ICA. The authors noted that most of the differences are explained by a change in blood vessel diameter. Accordingly, the sole examination of blood velocity may be insufficient to identify differences. However, these results from Sato et al. (2012) could also imply that regional differences in adaptation to reduction in blood pressure are only present extracranially, explaining the lack of difference in intracranial BV changes from the current study. Conversely, Ogoh et al. did not find any differences between ICA and VA blood flow changes during graded lower body negative pressure (Ogoh et al., 2015). Accordingly, the variability in cerebral blood flow/blood velocity changes in response to acute reduction in blood pressure, as well as the presence/absence of regional differences in these cerebrovascular responses, could also be explained by the chosen stimulus to manipulate blood pressure. Further research will be necessary to examine the influence of the different characteristics related to the stimulus used to acutely reduce blood pressure (e.g. graded vs. transient; active vs. passive) on regional cerebral blood flow/blood velocity changes.

### Lack of association between dCA and reduction in PCAv_mean_ induced by a sit-to-stand

We hypothesized that the posterior cerebral circulation would show a diminished dCA compared to MCA in these young fit women. This was based on women showing a greater prevalence of orthostatic symptoms (Ali et al., 2000). Since the posterior cerebral circulation irrigates cardiovascular and respiratory control centers, its hypoperfusion could cause the appearance of presyncopal symptoms. However, we did not find any correlations between PCAv_mean_ reduction upon standing and dCA metrics during the sit-to-stand procedure. Of note, studies comparing sex differences in orthostatic tolerance (Fu et al., 2004) and relating orthostatic tolerance to a greater PCAv_mean_ reduction (Kay et al., 2017) used a lower body negative pressure protocol. This method induces important and progressive decrease in MAP, whereas a sit-to-stand causes only an acute blood pressure drop, which is recovered within seconds. Therefore, the type of stimuli seems to be of importance when assessing regional differences in dCA and the impact of blood pressure reduction on PCAv_mean_ changes.

### TFA of forced oscillations in MAP and CBV

A very limited number of studies assessed dCA using TFA of forced oscillations in MAP and CBV in both MCA and PCA. Similarly to our results, the absolute gain was found to be greater in the MCA of young and healthy participants during 0.05 Hz and 0.10 Hz squat-stands (Smirl, Tzeng, Monteleone, & Ainslie, 2014). As previously discussed, this is caused by the MCA providing 70-75% of the blood to the brain compared to only 20-25% for the PCA. Since MCAv_mean_ and PCAv_mean_ are different at baseline, utilization of nGain to assess differences is more appropriate. Altogether, findings from our study and others (Smirl et al., 2014) demonstrate that TFA does not identify regional differences when dCA is evaluated with large and repeated MAP oscillations. Through the utilization of two different approaches to induce large MAP changes in order to characterize dCA, including the gold standard for linear TFA metric interpretation (Smirl et al., 2015), we believe these findings convincingly demonstrate the absence of difference in dCA between the MCA and PCA in these young fit women.

### Limitations

Limitations related to this study need to be acknowledged and further discussed. This study included only a small number of young healthy fit women and the results cannot be generalized to men or other populations (older individuals or patient populations). BP was measured by finger photoplethysmography. Even though we ensured participants avoided squeezing their fingers to optimize the signal, we acknowledge that BP may have been different compared to an invasive BP monitoring. Further to this point, MCAv_mean_ and PCAv_mean_ were monitored with transcranial Doppler ultrasound, and would be representative of flow only if the diameter of the arteries remains stable. Changes in P_ET_CO_2_ have been associated with changes in the diameter of the ICA and MCA. However, the physiological range of variation in P_ET_CO_2_ during all the tests in this study will most likely be associated with a minor effect on the diameter of the cerebral arteries (Coverdale, Gati, Opalevych, Perrotta, & Shoemaker, 2014; Verbree et al., 2014). Further research is thus needed to support our findings using invasive BP monitoring and volumetric flow measurements. Since female participants in this study were either taking oral contraceptives continuously (n=2), having an intrauterine device (n=2) or evaluated during days 1 to 10 of their menstrual cycle (n=7), we are unable to ascertain if dCA of both MCA and PCA was influenced by the oscillatory nature of ovarian hormones. Further research is warranted to determine the specific effects the stages of the menstrual cycle play on these measures.

### Conclusion

Using a multi-metrics approach including MAP oscillations of different natures and amplitudes, the results of this study indicate that dCA is comparable between the MCA and PCA in young fit women. In addition, cerebrovascular responses following a sit-to-stand are not different between these intracranial arteries and not associated with dCA metrics. Taken together, these findings indicate that dCA is not responsible for making the posterior cerebral circulation more vulnerable to transient reduction in blood pressure in these women.

## Author contribution

Data was collected at the Research center of the Institut universitaire de cardiologie et de pneumologie de Québec, Québec, Canada. P.B. was responsible for the conception and design of the work. L.L., K.R. and S.I. contributed to acquisition of data, L.L. analysed the data. All the authors contributed to the interpretation of data. L.L., A.D. and P.B. drafted the article. All authors provided approval of the final article, agree to be accountable for all aspects of the work in ensuring that questions related to the accuracy or integrity of any part of the work are appropriately investigated and resolved and that all persons designated as authors qualify for authorship, and all those who qualify for authorship are listed

